# UDP-Glucose 6-Dehydrogenase Knockout Impairs Migration and Decreases *in vivo* Metastatic Ability of Breast Cancer Cells

**DOI:** 10.1101/2020.05.30.125419

**Authors:** Shao Thing Teoh, Martin P. Ogrodzinski, Sophia Y. Lunt

## Abstract

Dysregulated metabolism is a hallmark of cancer that supports tumor growth and metastasis. One understudied aspect of cancer metabolism is altered nucleotide sugar biosynthesis, which drives aberrant cell surface glycosylation known to support various aspects of cancer cell behavior including migration and signaling. We examined clinical association of nucleotide sugar pathway gene expression and found that *UGDH*, encoding UDP-glucose 6-dehydrogenase which catalyzes production of UDP-glucuronate, is associated with worse breast cancer patient survival. Knocking out the mouse homolog *Ugdh* in highly-metastatic 6DT1 breast cancer cells impaired migration ability without affecting *in vitro* proliferation. Further, *Ugdh-KO* resulted in significantly decreased metastatic capacity *in vivo* when the cells were orthotopically injected in syngeneic mice. Our experiments show that UDP-glucuronate biosynthesis is critical for metastasis in a mouse model of breast cancer.

## INTRODUCTION

Breast cancer remains the most common cancer in women worldwide and is the leading cause of cancer deaths in women [1]. The vast majority of breast cancer-related deaths occur due to metastasis, which is currently considered incurable [2]. Deciphering the molecular events that support breast cancer growth and metastasis is therefore critical to developing new treatment strategies for combating this disease. Dysregulated metabolism is a hallmark of cancer that may present specific vulnerabilities for therapeutic targeting of cancer cells [3]. Metabolism is involved in more than mere production of energy and biosynthetic intermediates: metabolites play central roles in cell signaling [4–6], epigenetic regulation [7,8], and interaction with the microenvironment [9–11]. With the advent of mass spectrometry-based metabolite profiling technologies, metabolomics – the comprehensive and quantitative profiling of metabolites in biological systems – has emerged as a promising tool to reveal new insights about cancer metabolism and identify novel therapeutic targets.

One understudied aspect of cancer metabolism is aberrant nucleotide sugar biosynthesis. Nucleotide sugars are activated forms of monosaccharides, in which the high energy bond between the sugar moiety and the nucleotide allows the molecule to act as glycosyl donors in glycosylation reactions. Although nucleotide sugar biosynthetic pathways receive proportionally much smaller amounts of glucose flux compared to central metabolic pathways such as glycolysis and the tricarboxylic acid (TCA) cycle [12,13], recent studies have shown that metabolic flux through these pathways directly influences glycosylation by modifying nucleotide sugar availability [14–16]. Glycosylation of cell surface proteins mediate diverse aspects of cell behavior including cell-cell communication, adhesion, and migration [17]. Glycosylation of intracellular proteins is also important for mediating functions in signal transduction and gene regulation [18,19]. Other roles for nucleotide sugar-driven glycosylation are glycogen production for energy storage [20], and glucuronidation of xenobiotics to facilitate their metabolism and elimination [21]. Altered glycosylation is emerging as a common feature of cancers [17,22,23]; one recognized glycosylation feature of cancer cells is elevated presentation of N-acetylneuraminic acid (commonly known as sialic acid) on the cell surface [24]. We previously reported that sialic acid biosynthesis is elevated in highly-metastatic tumors and cancer cells relative to their less-metastatic counterparts, and that blockade of sialylation by deletion of a key gene in the sialic acid pathway leads to impaired *in vivo* metastatic ability in orthotopic tumor models [25].

Given the important role of glycosylation in mediating cancer cell behavior, it is likely that other nucleotide sugar metabolic pathways also support cancer metastasis. In addition to the sialic acid pathway, several nucleotide sugars – uridine diphosphate (UDP)-glucose, UDP-glucuronate and UDP-xylose – are produced via a parallel biosynthetic pathway (referred to as the ‘UDP-glucose pathway’ hereafter in this manuscript; **Figure 1**). In this study, we investigate the importance of the UDP-glucose pathway in breast cancer metastasis. We first determined the clinical relevance of this pathway based on correlation of gene expression with breast cancer patient outcomes. We found that a gene expression pattern which promotes UDP-glucuronate production and accumulation is correlated with decreased patient survival within poor-prognosis subsets of breast cancer patients. We then knocked out *Ugdh*, the rate-limiting gene mediating UDP-glucuronate production, in 6DT1 mouse breast cancer cells and found that *Ugdh* knockout impaired motility of these highly-metastatic cells. We further show that *Ugdh* knockout impairs *in vivo* tumor growth and metastasis, demonstrating the importance of this gene in breast cancer progression. Our results suggest that attenuation of UDP-glucuronate production by targeting *UGDH* is a potential strategy to inhibit breast cancer metastasis.

**Figure 1.**
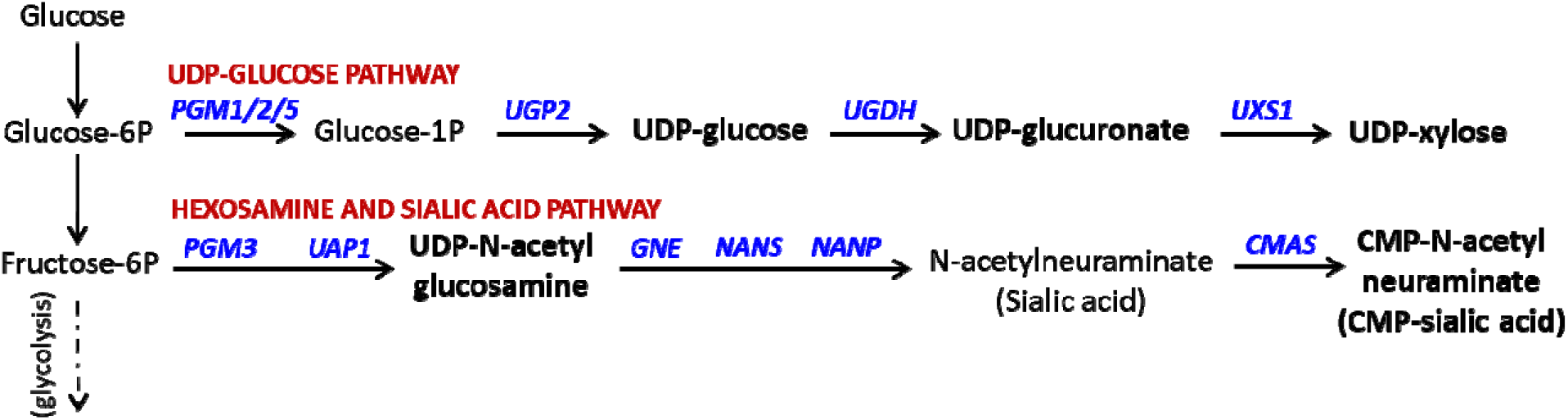
Two parallel nucleotide sugar-producing pathways with significance in breast cancer. Nucleotide sugars are highlighted in bold.

## RESULTS

### High *UGDH* expression is associated with decreased patient survival in poor-prognosis breast cancer subsets

To examine correlation between high expression of genes in the UDP-glucose pathway and patient survival, we used KM Plotter [26], an online tool that interrogates gene expression and patient survival data in publicly-available Gene Expression Omnibus (GEO) datasets [27] (**Figure 2**). We found that across all breast cancer cases, high *UGP2* expression correlates with decreased patient survival (hazard ratio (HR) = 1.21, p = 0.00056), but other pathway genes *UGDH* and *UXS1* are not clearly associated with differences in patient survival (HR = 0.9 and 0.97, respectively; **Figure 2A**). Since breast cancer is heterogeneous and different subtypes of breast cancer vary in their metabolic characteristics [28], we analyzed specific subtypes of breast cancer. We first stratified the patient data by the four intrinsic molecular subtypes – luminal A, luminal B, HER2-enriched and basal, which are classified according to gene expression profile [29]. We found that *UGP2* and *UGDH* expression is correlated with significantly worse patient survival (HR = 1.38, p = 0.021 and HR = 1.47; p = 0.0098) in the basal subtype (**Figure 2B**), which has the worst prognosis [30,31], but not the other subtypes (**Supplementary Figure 1**).

**Figure 2.**
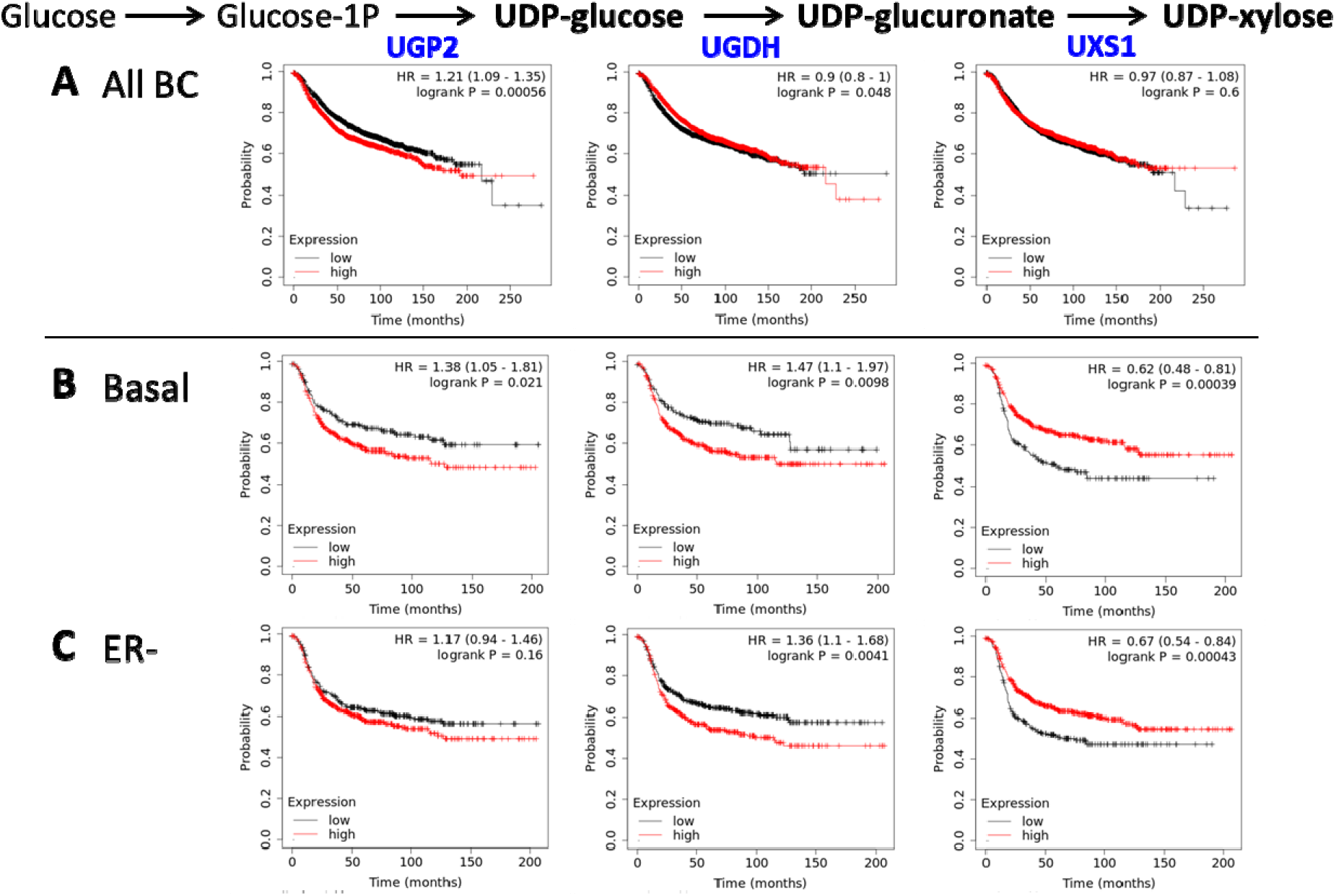
High UGDH and low UXS1 expression correlates with worse patient survival in poor-prognosis subsets of breast cancer. Kaplan-Meier survival curves showing relapse-free survival for patients with high and low expression of genes in the UDP-glucose pathway, generated in KM Plotter for (A) all breast cancer, (B) basal intrinsic subtype breast cancer, which has the worst prognosis among intrinsic subtypes; and (C) ER-negative breast cancer, which has worse prognosis than ER-positive breast cancer. Cut-off values for splitting high-versus low-expression patient groups were automatically determined using the ‘Auto select cutoff’ option in KM Plotter. Hazard ratios and p-values are also listed in **Table 1**.

We also stratified the patient data by estrogen receptor (ER) status and found that *UGDH* expression is correlated with significantly worse patient survival (HR = 1.36, p = 0.0041) in ER-negative (ER-) breast cancer (**Figure 2C**), but not in ER-positive (ER+) breast cancer (**Supplementary Figure 2**). ER-breast cancer generally has worse prognosis than ER+ breast cancer [32–34]. Thus, high expression of *UGP2* and *UGDH* correlate with worse patient survival specifically within subsets of breast cancer with poor prognosis. Interestingly, in both the basal and ER-subsets, high *UXS1* expression correlated with *increased* patient survival – or, conversely, low *UXS1* expression correlated with decreased patient survival (**Figure 2B-C**). In the UDP-glucose pathway, *UGP2* and *UGDH* are both upstream of UDP-glucuronate, while *UXS1* is downstream (**Figure 1**). High *UGP2* and *UGDH* expression would increase UDP-glucoronate production while low *UXS1* would decrease UDP-glucuronate consumption; both have the net effect of increasing UDP-glucuronate levels. The results (summarized in **Table 1**) therefore suggest that increased UDP-glucuronate levels may be linked with decreased patient survival within poor-prognosis breast cancer subsets.

**Table 1.** Hazard Ratio (HR) and p-values (p) for high vs low expression of UDP-glucose pathway genes in various subsets of breast cancer patients. HR values larger than 1 indicate that the high-expression cohort has shorter survival times, while HR values smaller than 1 indicate that the high-expression cohort has longer survival times. Values in parentheses indicate the 95% confidence range. HR values with p < 0.05 are shown in bold. HR values with p < 0.01 are shaded in red (HR > 1; significant correlation with worse survival) or green (HR < 1; significant correlation with better survival). BC, breast cancer; ER, estrogen receptor; HER2, human epidermal growth factor 2.

| Subtype | Prognosis | UGP2 |  | UGDH |  | UXS1 |  |
| --- | --- | --- | --- | --- | --- | --- | --- |
|  |  | HR | p | HR | p | HR | p |
| all BC |  | **1.21 (1.09-1.35)** | **0.00056** | **0.9 (0.8-1)** | **0.048** | 0.97 (0.87-1.08) | 0.6 |
| luminal A | best | 1.21 (1-1.48) | 0.054 | 0.83 (0.69-1) | 0.05 | **0.77 (0.65-0.92)** | **3.7E-3** |
| luminal B | good | **1.3 (1.06-1.58)** | **0.01** | 1.18 (0.97-1.44) | 0.094 | **0.8 (0.66-0.97)** | **0.026** |
| HER2-enriched | poor | 1.94 (0.98-3.85) | 0.053 | 1.95 (0.97-3.93) | 0.058 | 1.67 (0.88-3.18) | 0.11 |
| basal | worst | **1.38 (1.05-1.81)** | **0.021** | **1.47 (1.1-1.97)** | **9.8E-3** | **0.62 (0.48-0.81)** | **3.9E-4** |
| ER+ | good | **1.22 (1.06-1.41)** | **6.4E-3** | 0.89 (0.78-1.02) | 0.1 | **0.85 (0.74-0.97)** | **0.018** |
| ER- | poor | 1.17 (0.94-1.46) | 0.16 | **1.36 (1.1-1.68)** | **4.1E-3** | **0.67 (0.54-0.84)** | **4.3E-4** |

### Ugdh KO decreases migratory ability of breast cancer cells

Given the clinical significance highlighted by KM Plotter, we investigated whether UDP-glucuronate production is essential to breast cancer cells. To evaluate this, we performed CRISPR/Cas9-mediated knockout (KO) of *Ugdh* (the mouse homolog of human *UGDH*) in the highly-metastatic 6DT1 mouse mammary cancer cell line. The 6DT1 cell line was chosen for our model because the orthotopic tumors generated are triple negative by immunohistochemical subtyping, have a gene expression signature characteristic of claudin-low breast cancer (a poor prognosis breast cancer subtype with stem cell-like features), and transcriptionally co-cluster with basal breast cancer patient derived xenografts [35]. 6DT1 cells were transiently transfected with ribonucleoprotein (RNP) complexes of Cas9 ribonuclease and *Ugdh*-targeting small guide RNA, and a clonal population of *Ugdh* knockout (*Ugdh*-KO) cells with confirmed frame-shifted *Ugdh* gene sequence (**Supplementary Figure 3**) was isolated. In parallel, a control population with wild-type *Ugdh* was isolated in the same transfection experiment and used as wild-type (WT) controls. We found that *Ugdh-KO* and WT cells have similar *in vitro* proliferation rates (**Figure 3A**). We next examined the impact of *Ugdh* KO on migration ability. First, we performed a modified scratch assay to measure wound-healing ability [36]. We used silicone well inserts to generate migration gaps to avoid scratching the plate surface, which causes cell damage and death adjacent to the scratch and impacts migration [37]. We found that *Ugdh-KO* cells take significantly longer to close the gap compared to WT cells, indicating loss of migration ability (**Figure 3B**). We also measured chemotactic migration using the Boyden chamber (also known as Transwell) assay [38]. We found that far fewer *Ugdh-KO* cells successfully migrate across a cell-permeable membrane towards the nutrient gradient (**Figure 3C**, **Supplementary Figure 4**), confirming that migration is indeed impaired in *Ugdh-KO* cells.

**Figure 3.**
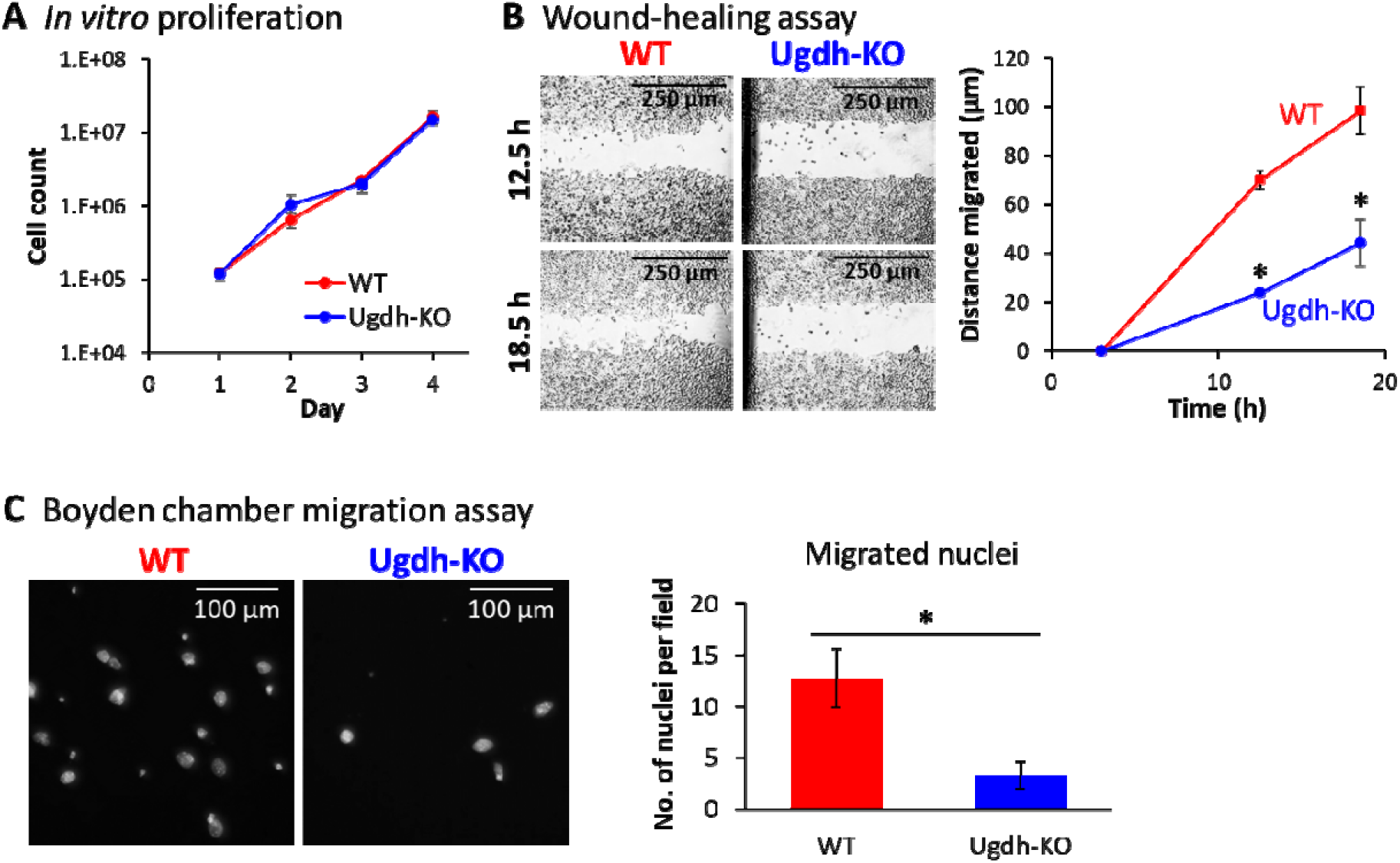
Ugdh-KO cancer cells have decreased migratory ability. (A) Proliferation curve showing no difference in WT versus Ugdh-KO cell counts over 4 days. The vertical axis shows cell count on a logarithmic scale. Values are the average of 3 measurements and error bars are standard deviations. (B) Measurement of migration ability by wound healing (gap closure) assay shows a significant difference between WT and Ugdh-KO cells. Left: Representative images of WT and Ugdh-KO cells at 12.5 and 18.5 h after starting the wound healing assay. Right: quantification results of migrated distance, relative to start point at t = 3 h. Values are averages of 6 images (2 locations per well, 3 well replicates). (C) Measurement of chemotactic migration by the Boyden chamber assay shows a significant difference between WT and Ugdh-KO cells. Left: Representative fluorescent images of Hoechst 33342-stained cells on underside of Transwell inserts, showing large, oval-shaped nuclei of fully migrated cells. Right: barplot showing number of fully migrated nuclei following 16 hours of migration. Barplot values and error bars are averages and standard deviations, respectively, of 27 images (9 images per well, 3 well replicates). Asterisks (*) denote statistical significance (p-value < 0.05 by Welch’s t-test) in the difference between WT and *Ugdh-KO*.

### *Ugdh* KO blocks UDP-glucuronate production

To investigate the metabolic effects of *Ugdh* KO, we next performed a metabolomics comparison of *Ugdh-KO* and WT cells using liquid chromatography-mass spectrometry [25]. *Ugdh* KO abolished UDP-glucuronate production, confirming that *Ugdh* is the sole enzyme catalyzing UDP-glucuronate production; we also observed slightly increased accumulation of upstream UDP-glucose due to the blockade at *Ugdh* (**Figure 4A**). Interestingly, UDP-xylose, the downstream metabolite of UDP-glucuronate, was only diminished by ~50% compared to WT (**Figure 4A**), suggesting the presence of an alternate route for UDP-xylose biosynthesis. Several other metabolite peaks showed significantly altered peak intensities in *Ugdh-KO:* GAP+DHAP, UMP and N-acetylmannosamine-6-phosphate (**Figure 4B**). The combined level of glyceraldehyde 3-phosphate and dihydroxyacetone phosphate (GAP+DHAP), main glycolytic intermediates, was increased over 2-fold in *Ugdh-KO*, likely due to carbon flux in the blocked UDP-glucose pathway overflowing into glycolysis. Similarly, increased N-acetylmannosamine-6-phosphate is likely due to overflow from UDP-glucose pathway resulting in increased flow of carbon into the hexosamine/sialic acid pathway. This is corroborated by slightly increased levels of all other hexosamine/sialic acid pathway metabolites: N-acetylglucosamine-1-phosphate, UDP-N-acetylglucosamine, N-acetylneuraminate, and CMP-N-acetylneuraminate (**Supplementary Figure 5A**). On the other hand, uridine monophosphate (UMP) is decreased in *Ugdh-KO;* this may reflect decreased pyrimidine biosynthesis or decreased cycling of uridine triphosphate (UTP) to UMP via generation of UDP-sugars. Besides the above, there were few other significant metabolite changes, indicating that disruption of this pathway did not globally perturb cellular metabolism (**Supplementary Figure 5B**).

**Figure 4.**
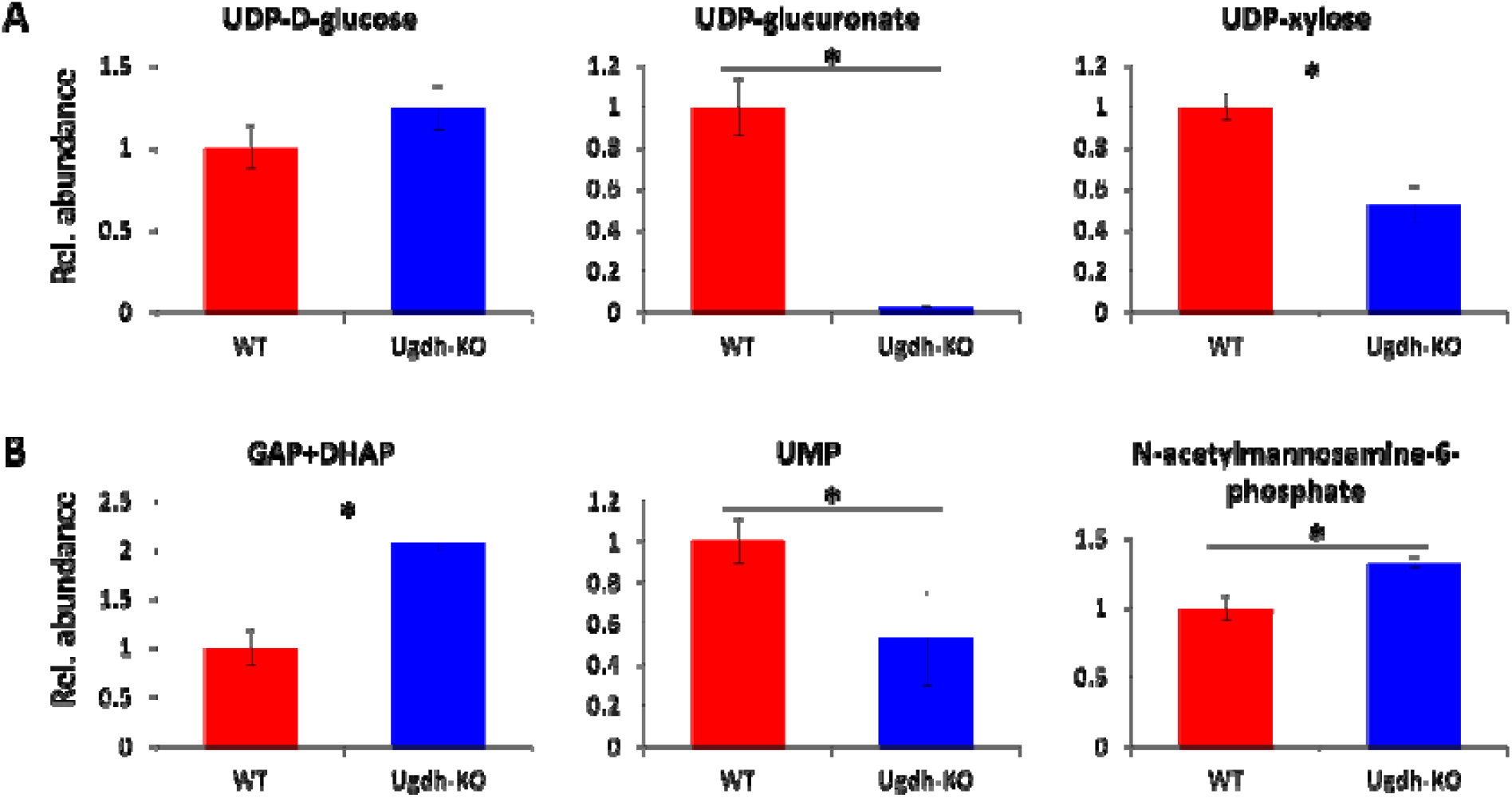
Ugdh-KO abolishes UDP-glucuronate but not UDP-xylose production. Relative abundance of (A) metabolites in the UDP-glucose pathway, (B) other metabolites significantly altered in Ugdh-KO. Values are displayed relative to wild-type (WT) cells and represent average of 3 replicates. Error bars represent standard deviations. Asterisks (*) denote statistical significance (p-value < 0.05 by Welch’s t-test).

### *Ugdh* KO does not impair epithelial-mesenchymal transition (EMT)

A recent study found that *UGDH* KO in human lung cancer cells resulted in impaired epithelial-mesenchymal transition (EMT), which impacted cell migration and extravasation [39]. To examine whether EMT is similarly impaired by *Ugdh* KO in our mouse breast cancer cells, we quantified the gene expression levels of a panel of EMT-associated transcription factors (*Elf5, Foxc2, Grhl2, Six1, Snai1, Snai2, Tcf3, Tcf4, Twist1, Zeb1, Zeb2*) and markers (*Cdh1, Cdh2, Fn1, Vim*). We found three genes significantly upregulated in *Ugdh-KO* cells: *Cdh1*, encoding the epithelial marker E-cadherin; *Fn1*, encoding the extracellular glycoprotein fibronectin; and *Six1*, a transcription factor implicated in proliferation and metastasis of tumor cells (**Figure 5**). EMT is associated with downregulation of *Cdh1* and upregulation of *Cdh2* (encoding N-cadherin, a mesenchymal cell marker). Although increased expression of *Cdh1* is partially consistent with decreased EMT, *Cdh2* expression was unchanged in *Ugdh-KO*. Additionally, fibronectin is a recognized mesenchymal cell marker [40,41], and the *Six1* gene product is associated with a role in promoting EMT [42–44]. However, *Fn1* and *Six1* expression are both increased, rather than decreased, in *Ugdh-KO* cells. Therefore, our data shows that EMT is not transcriptionally inhibited by *Ugdh* KO, suggesting that impaired EMT is unlikely to be the cause of decreased migration in these cells.

**Figure 5.**
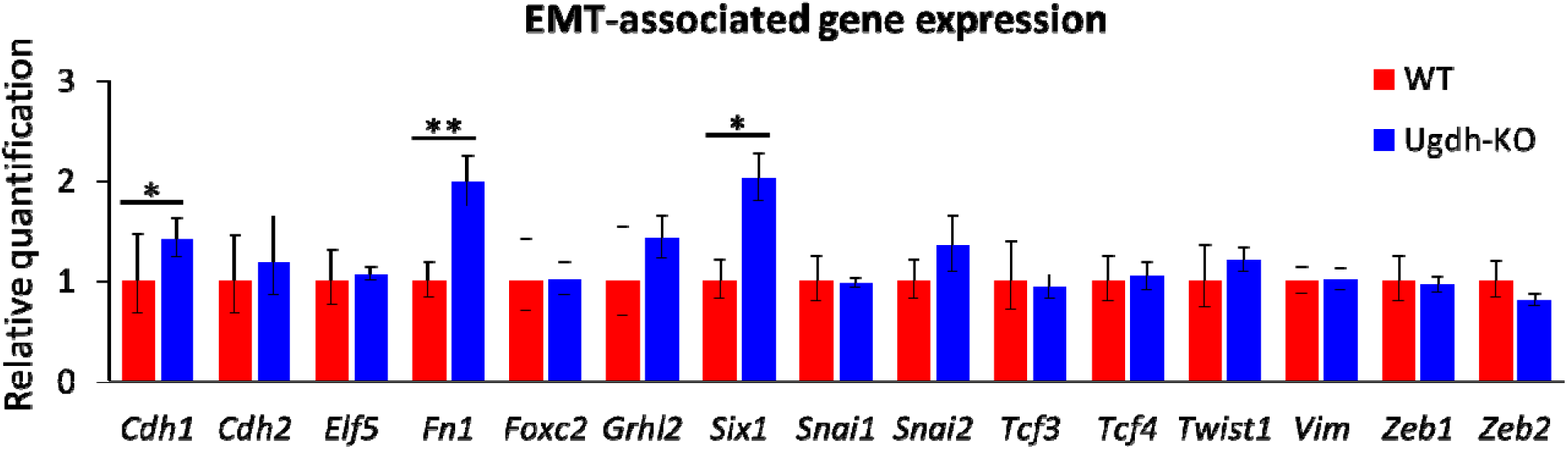
*Ugdh-KO* does not transcriptionally inhibit EMT in 6DT1 cells. Expression levels of genes associated with EMT in 6DT1 cells quantified by qPCR. Only *Cdh1, Fn1* and *Six1* are transcriptionally upregulated in *Ugdh-KO* cells; of these, only *Cdh1* upregulation is consistent with decreased EMT. All expression levels are normalized to control gene Tubb5 and displayed relative to WT. Relative quantitation values shown are averages of 4 replicates (2 cell culture replicates × 2 PCR plate replicates). Error bars represent ranges in the relative quantitation values, calculated from standard deviation of corresponding. ΔΔC_T_ values (relative quantitation, RQ = 2^ΔΔC_T_). Statistically significant differences between WT and Ugdh-KO are indicated by asterisks: *, p < 0.05 and **, p < 0.01, where p-values are calculated from ΔΔC_T_ values using Welch’s t-test. ΔΔCT: difference in *real-time PCR cycle threshold difference between gene of interest and control gene* between Ugdh-KO and WT samples.

### *Ugdh* KO decreases tumor growth and lung metastasis *in vivo*

Finally, to evaluate the *in vivo* effects of UDP-glucuronate depletion, we orthotopically injected 6DT1 *Ugdh-KO* or WT cells into syngeneic FVB mice. Both *Ugdh-KO* and WT injections produced tumors that were detectable within 7 days post injection; however, *Ugdh-KO* tumors grew noticeably slower. At 28 days post injection, several mice in the WT-injected cohort began to exhibit labored breathing and decreased activity; we therefore sacrificed both mouse cohorts and examined them for pulmonary metastases. In general, *Ugdh-KO* tumors were smaller than WT tumors as evidenced by lower tumor weights at endpoint (**Figure 6A**), although the difference lacked statistical significance (p = 0.08) due to relatively large variance between individual samples. The effect of *Ugdh* KO on the extent of lung metastasis was much more significant: while WT-injected mice had numerous gross metastatic lesions throughout the lungs, consistent with the high metastatic capacity of the 6DT1 cell line, mice injected with *Ugdh-KO* cells were virtually free of lung metastases (**Figure 6B**). To gain a quantitative representation of the extent of lung metastasis, we measured the area of metastatic tissue relative to total lung area (**Figure 6C**), as well as the number of discrete metastatic lesions (**Figure 6D**). In both cases, the extent of lung metastasis was significantly lower in *Ugdh-KO* injected mice compared to WT injected mice. To verify that the effects of *Ugdh* KO is due to loss of UDP-glucuronate and not its downstream metabolite UDP-xylose (**Figure 1**), we additionally generated *Uxs1-KO* cells. We confirmed that *Uxs1*-KO resulted in greatly reduced UDP-xylose production (**Figure 7A**) and accumulation of UDP-glucuronate (**Figure 7B**), but did not affect tumor growth or lung metastasis (**Figure 7C-F**). Thus, depletion of UDP-glucuronate by *Ugdh* KO in highly-metastatic 6DT1 cells results in decreased tumor growth and lung metastatic ability.

**Figure 6.**
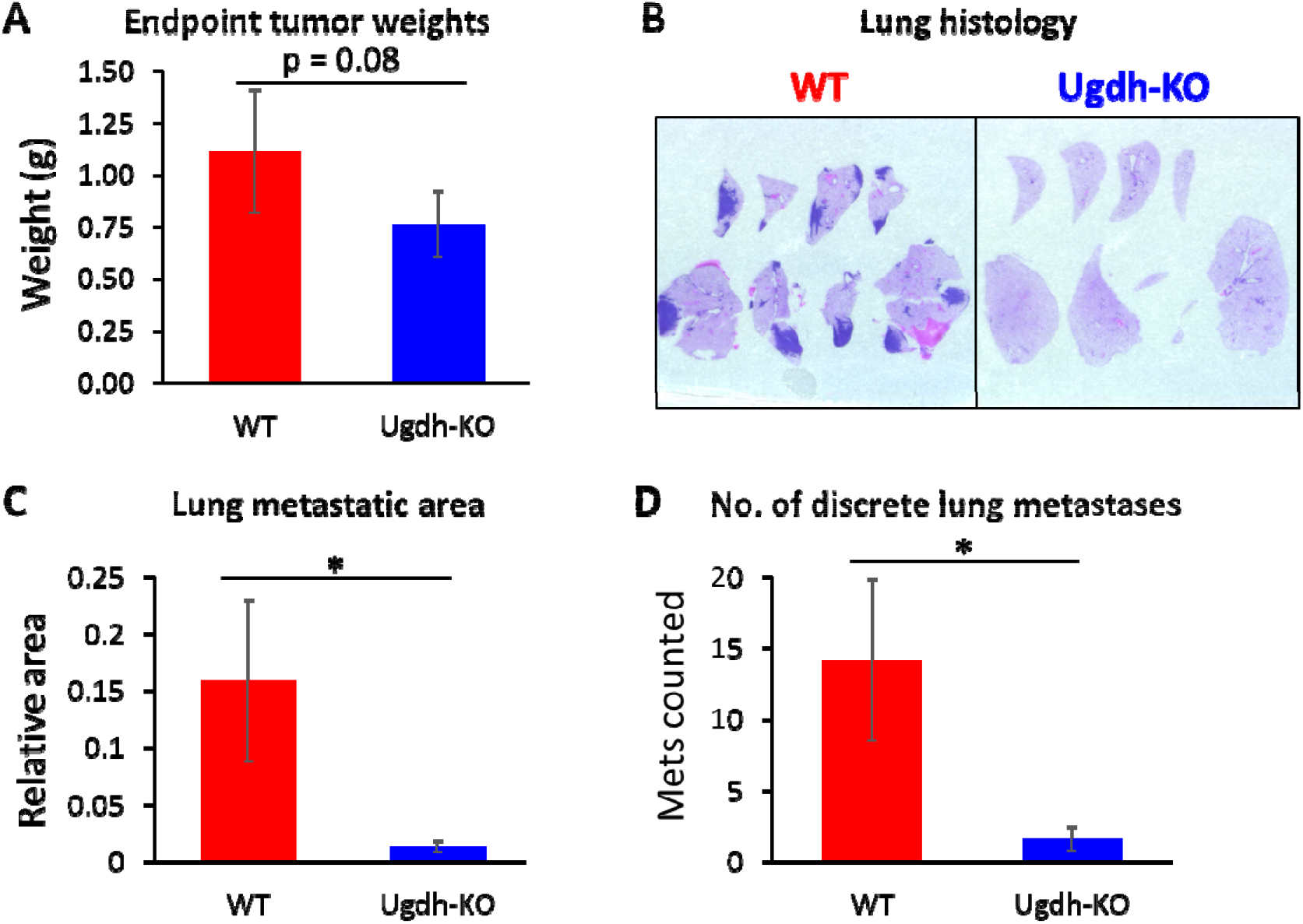
Ugdh-KO cells grow and metastasize poorly following orthotopic implantation. (A) Tumor weights at endpoint. (B) Representative images of hematoxylin and eosin (H&E) stained lung sections from mice injected with WT or Ugdh-KO cells. Dark blue regions are metastatic tumor tissue, which are nuclei-dense and heavily stained by blue hematoxylin dye. (C) Metastatic tissue area relative to total lung area in mice injected with WT or Ugdh-KO cells. (D) Number of discrete metastatic lesions counted on lung sections of mice injected with WT or Ugdh-KO cells. All values represent averages of 5 animal replicates. Error bars represent SD. Statistically significant differences (p-value < 0.05) are marked with asterisks (*).

**Figure 7.**
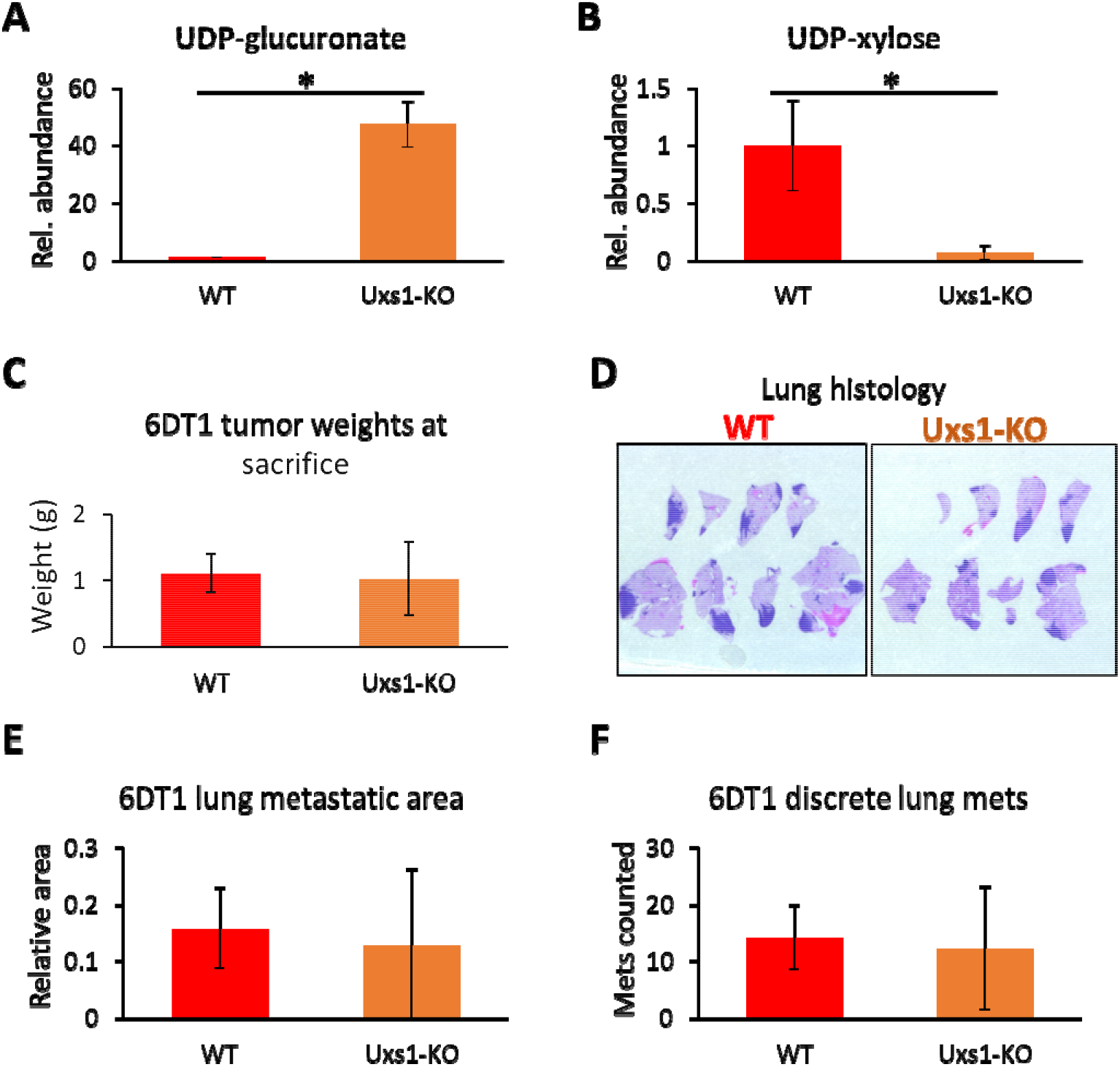
Uxs1 KO does not significantly affect tumor growth or metastasis. (A) Relative abundance of UDP-xylose in WT, Ugdh-KO and Uxs1-KO cells. Values are displayed relative to WT and represent average of 3 replicates. Error bars represent standard deviations. Asterisks (*) denote statistically significant difference (p-value < 0.05 by Welch’s t-test) compared to WT. (B) Tumor weights at endpoint of 28 days post-injection. (C) Representative images of hematoxylin and eosin (H&E) stained lung sections from mice injected with WT vs Uxs1-KO cells. Dark blue regions are metastatic tumor tissue, which are nuclei-dense and heavily stained by blue hematoxylin dye. (D) Metastatic tissue area relative to total lung area in mice injected with WT or Uxs1-KO cells. (E) Number of discrete metastatic lesions counted on lung sections of mice injected with WT or Uxs1-KO cells. Values presented in B, D and E are averages of 5 animal replicates and error bars represent standard deviation.

## DISCUSSION

Our experiments show that *Ugdh* KO depletes intracellular UDP-glucuronate levels and decreases tumor growth and metastatic ability of highly metastatic breast cancer cells. Notably, we show that migration ability is severely impacted in *Ugdh-KO* cells according to both the wound healing and Boyden chamber assays (**Figure 3B-C**). This loss of migration ability likely contributed to the reduction in metastatic ability, since migration and invasion through surrounding tissues is a key step in the metastatic process [45,46]. Another study also found that *UGDH* KO impairs migration and lung colonization ability in human lung cancer cells [39]. Interestingly, the authors showed that it was accumulation of UDP-glucose in *UGDH-KO* cells – rather than depletion of UDP-glucuronate – that led to destabilized *SNAI1* mRNA, impaired EMT, decreased migration ability and ultimately resulted in loss of extravasation ability required for circulating tumor cells to colonize the lung. In contrast, our findings did not indicate reduced expression of EMT-associated genes in *Ugdh-KO* 6DT1 breast cancer cells (**Figure 5**). We additionally examined *Snai1* mRNA stability in our cells and determined that there was no difference between *Ugdh-KO* and WT (**Supplementary Figure 6**). One possible reason for this discrepancy is that our cells do not accumulate UDP-glucose to significantly higher levels upon *Ugdh* KO (**Figure 4**), possibly because excess UDP-glucose is converted to glycogen or used for protein glycosylation more efficiently in these cells. Therefore, the reduction in tumor growth and metastasis in our model likely proceeds through different mechanisms. In turn, this suggests that *UGDH* is a strong candidate for pharmacological targeting, since its inhibition may have multiple effects that potentially impact tumor growth and metastasis.

In our qPCR data, the EMT-associated gene that changed most significantly in *Ugdh-KO* was *Fn1*, which encodes the extracellular glycoprotein fibronectin (**Figure 5**). Fibronectin is a mesenchymal cell marker that plays major roles in cell adhesion, growth, migration, and differentiation, and it is important for processes such as wound healing and embryonic development [40,41]. The presence of fibronectin in the extracellular matrix has been shown to induce EMT in breast cancer and normal mammary cells [47,48]. It is therefore possible that in breast cancer cells, increased *Fn1*/fibronectin expression may act to maintain EMT, which may explain why EMT was not impaired in our *Ugdh-KO* cells as observed elsewhere [39]. Interestingly, it has been reported that removal of hyaluronic acid (a major downstream product of UDP-glucuronate) from the extracellular matrix – either by inhibition of hyaluronic acid synthesis or digestion of matrix hyaluronic acid by hyaluronidase – significantly increased fibronectin gene expression and extracellular fibronectin protein deposition [49]. Since Ugdh-KO abolishes UDP-glucuronate and prevents hyaluronic acid synthesis, it is possible that the same regulatory mechanism is responsible for increasing *Fn1* expression in our *Ugdh-KO* cells.

In our metabolomics comparison of *Ugdh-KO* vs WT cells, we found a diminished but surprisingly substantial level of UDP-xylose remaining in *Ugdh-KO* cells despite UDP-glucuronate production being completely abolished (**Figure 4**). This suggested that UDP-xylose may not be produced solely from UDP-glucuronate. In fact, an alternate pathway involving broad substrate UDP-sugar pyrophosphorylase (USP) has been characterized in plants [50] as well as the protozoan *Leishmania major* [51]. A potentially related finding was elevation of xylulose-5-phosphate, which is a close structural isomer to xylose phosphate, in *Ugdh-KO* (**Supplementary Figure 5C**). Given that the levels of two other intermediates in the pentose phosphate (PPP), ribulose-5-phosphate and ribose-5-phosphate, did not noticeably change (**Supplementary Figure 5C**), it is possible that an alternate pathway to UDP-xylose exists via conversion from xylulose-5-phosphate, which may be upregulated in compensation for UDP-glucuronate shortage. However, a broad substrate USP has not been identified in mammalian cells, and further investigation of this potential alternate pathway is beyond the scope of this work.

In conclusion, we demonstrated that *Ugdh-KO* in breast cancer cells greatly reduces *in vivo* metastasis through attenuation of migration ability. Together with a previous report in lung cancer [39], this establishes a functional role of *Ugdh* in cancer metastasis. Our results highlight the importance of dysregulated nucleotide sugar metabolism in breast cancer metastasis and underscore the need to develop targeted therapeutics that can inhibit these metabolic pathways.

## METHODS

### Survival Analysis

Survival curves were generated using KM Plotter for Breast Cancer [26] using probes 205480_s_at for *UGP2*, 203343_at for *UGDH* and 219675_s_at for *UXS1*. Patients were split into high and low expression groups using the ‘Auto select best cutoff’ option. Redundant samples were removed and biased arrays were excluded as per the default quality control settings.

### Cell culture

6DT1 mouse mammary cancer cells were cultured in Dulbecco’s Modified Eagle Medium (DMEM Corning, Corning, New York 10-017-CM) with 25 mM glucose without sodium pyruvate supplemented with 2 mM glutamine (Corning, 25-005-CI) 10% heat-inactivated fetal bovine serum (MilliporeSigma, Burlington Massachusetts, 12306C), and 1% penicillin and streptomycin (Corning, 30-002-CI). Cells were maintained at 37°C with 5% CO2. For proliferation assays, cells were seeded at a density of 50,000 cells/well in 6-well tissue culture plates then counted daily for 4 days using a Nexcelom Cellometer Auto T4 cell counter.

### CRISPR/Cas9

Single-stranded guide RNAs (sgRNAs) targeting exon 2 of mouse *Ugdh* and exon 3 of mouse *Uxs1* were designed using Benchling: 5’-TACGGTTGTGGATGTCAACG (Ugdh) and 5’-GGGGCAACTTTGTTAACATG (Uxs1). CRISPR/Cas9 genome editing was performed using the Alt-R^®^ CRISPR-Cas9 System (Integrated DNA Technologies) as previously described [25]. Briefly, Alt-R CRISPR-Cas9 cRNA containing Cmas target sequence, ATTO 550 fluorophore-linked Alt-R CRISPR-Cas9 tracRNA and Alt-R S/p Cas9 Nuclease 3NLS were combined to form ribonucleoprotein (RNP) complexes targeting Ugdh or Uxs1. RNPs were delivered to the cells by reverse transfection using Lipofectamine RNAiMax (Thermo Fisher). 48 h after transfection, cells were trypsinized, counted, and resuspended to a concentration of 1 cell per 200 μL and seeded on 96-well plates at 200 μL/well. Plates were monitored for formation of single colonies indicating clonal populations derived from single cells. Once confluent, clonal populations were passaged to 24-well plates and expanded. Genomic DNA was extracted using DNeasy Blood and Tissue Kit (Qiagen) to check for successful integration of indels into the target region. PCR primers were designed to amplify a region spanning the sgRNA target site: forward 5’-TCCTTTAACCCACGTCCACCTG, reverse 5’-ACACCAGGCAGACTTTGGACTT (Ugdh); forward 5’-TGCTCAGGCACCTGGTTGATAA, reverse 5’-TCCCTGAGGAAAAAGGCAACCA (Uxs1). PCR products were agarose gel-purified, sequenced by ACGT Inc and the results analyzed using Tracking of Indels by Decomposition (TIDE) [52] to identify clones harboring indels in the target genes.

### Migration assays

For the wound healing assay, 2-chamber silicone inserts (Ibidi) were placed in 24-well plates using sterile tweezers. Cells were suspended at a concentration of 1×10^6^ cells/mL, then 70 μL of the cell suspension was added to each chamber of the inserts and grown overnight. The next day, inserts were removed using sterile tweezers to expose the migration gap, then media was aspirated and cells gently washed 3 times with PBS, and 500 μL DMEM containing 10 μg/mL Mitomycin C was added to each well. Each migration gap is marked in two locations by perpendicular lines drawn on the bottom surface of plate. At each of these locations, images are taken 3 h, 12.5 h and 18.5 h after starting the migration. The gap width was manually measured in each image using the ‘Measure distance between two perpendicular lines’ feature of the AmScope microscopy software. The 3 h time point was defined as the start point, and migrated distances were calculated from the difference in gap distance between the start point and subsequent time points. Values shown are averages of 6 images (2 locations per well, 3 well replicates).

For the Boyden chamber assay, tissue culture plate inserts with 8.0 μm pore size polycarbonate membranes (VWR) were placed in 24-well tissue culture plates (VWR), and 500 μL DMEM added to the lower chamber of each well. Cells were pretreated with 10 μg/mL Mitomycin C for 2 hours, then trypsinized and resuspended at 500,000 cells/mL in serum-free DMEM, and 100 μL was added to the upper chamber of the inserts. Cells were allowed to migrate for 16 h, then inserts were removed, media aspirated from the upper chamber, and cells remaining in the upper chamber were removed with cotton swabs. Inserts were placed in new 24-well plates containing 500 μL formalin in the lower chamber for 15 min to fix the cells attached on the bottom surface of the membranes. Inserts with fixed cells are washed 3 times by dipping in additional wells filled with PBS. Cells were then stained with 2 μM Hoechst 33342 for 5 min, washed twice with PBS, then imaged with a Leica DFC9000GT fluorescence camera. For each insert membrane, images were taken at 9 random locations in a roughly 3 x 3 grid pattern. Small, circular nuclei corresponding to the shape and size of the pores were considered to represent partially migrated cells (cells in the process of migrating) while larger, oval-shaped nuclei were considered fully migrated cells that spread out horizontally across the bottom membrane surface after passing through the pores. Nuclei were counted using ImageJ. Values shown are averages of 27 images (9 images per well, 3 well replicates).

### Metabolic profiling

Targeted metabolite profiling was performed as previously described [25,53]. Briefly, cells were seeded in 6-well tissue culture plates at 150,000 cells/well and cultured for 24 hours. Cells were washed with saline (VWR, Radnor, Pennsylvania, 16005-092) and metabolism was quenched with addition of cold methanol, followed by water containing 1 μM camphorsulfonic acid as internal standard. Extracts were then transferred to 1.5 mL Eppendorf tubes and cold chloroform was added to each tube and vortexed for 10 minutes at 4°C. The final metabolite extraction solvent ratios were methanol:water:chloroform (5:2:5). The polar phase was collected and dried under a stream of nitrogen gas. The dried metabolites were then resuspended in HPLC-grade water for analysis. For analysis of amino acids, a 20 μL aliquot of the sample was added to 80 μL MeOH and derivatized with 10 μL triethylamine and 2 μL benzylchloroformate. LC-MS/MS analysis was performed with ion-pairing reverse phase chromatography using an Ascentis Express column (C18, 5 cm x 2.1 mm, 2.7 μm, MilliporeSigma, 53822-U) and a Waters Xevo TQ-S triple quadrupole mass spectrometer. Mass spectra were acquired using negative mode electrospray ionization operating in multiple reaction monitoring (MRM) mode. Peak processing was performed using MAVEN [54]. For each sample, data from the non-derivatized and CBZ-derivatized sample runs were separately scaled to internal standard (camphorsulfonic acid) then combined, and the entire dataset normalized by Probabilistic Quotient Normalization [55]. Heatmaps were generated using Cluster 3.0 [56] and exported using Java Treeview [57].

### Statistical analyses

The nonparametric log-rank test was used for testing significance of difference between Kaplan-Meier survival curves. All other statistical analyses were performed using Welch’s t-test. The significance threshold was set at p = 0.05 for all statistical analyses. All error bars presented represent standard deviation.

### *In vivo* tumor studies

All animal use was performed in accordance with institutional and federal guidelines. 100,000 6DT1 cells were inoculated into the fourth mammary fat pad of 7- to 9-week-old FVB female mice. Mice were euthanized 28 days after injection by carbon dioxide asphyxiation and cervical dislocation. Lungs were harvested, fixed in formalin, embedded in paraffin, sectioned and stained with hematoxylin and eosin (H&E) for histology analysis.

### Histological analysis of lung metastasis

Microscope images of lung histology slides were acquired with a Leica M165FC stereo microscope. Image processing was performed using the FIJI distribution of ImageJ (version 1.52p). The color images were first deconvoluted into H (hematoxylin) and E (eosin) color channels using Color Deconvolution (‘H&E hide’ deconvolution matrix). Deconvoluted H and E images were saved as new TIFF images. To quantitate total lung area, E images were opened, smoothing was applied 10 times, automatic thresholding was applied to separate lung tissue from background, and the total area of thresholded regions was quantified. To quantitate lung metastasis, H images were opened, Auto Local Threshold was performed using Phansalkar’s algorithm with default parameters and a radius setting of 15. Total area of thresholded regions was quantified to obtain lung metastatic area, and number of discrete regions was enumerated to count the discrete metastases.

### qRT-PCR

Cells were seeded in 6-well tissue culture plates at 150,000 cells/well and cultured for 24 hours. Total RNA was extracted from the cells using the RNeasy Mini Kit (Qiagen, 74104), on-column DNase digestion performed using DNase I (Qiagen, 79254). cDNA was prepared using LunaScript^™^ RT SuperMix Kit (New England Biolabs, E3010S). Real-time PCR was performed using Luna^®^ Universal qPCR Master Mix (New England Biolabs, M3003S) on an Applied Biosystems StepOnePlus^™^ Real-Time PCR system with the following conditions: 10 min at 55 °C, 1 min at 95 °C followed by 40 cycles at 95 °C for 10 s and 60 °C for 1 min. Gene expression values were normalized to control gene Tubb5 and verified against an additional control gene Actb. The primer sequences used for real-time PCR are listed in Supplementary Table 1.

## AUTHOR CONTRIBUTIONS

ST performed gene knockout experiments, migration assays, metabolomics experiments, histology image analysis, qRT-PCR experiments, and data analysis. MO maintained the FVB mice used in the experiments and performed orthotopic cell injections and mouse necropsies. SL designed and supervised the study. All authors contributed to writing, reviewing, and/or revising the manuscript.

## ACKNOWLEDGMENTS

The authors thank Deanna Broadwater, Elliot Ensink, and Hyllana Medeiros for helpful discussions and critical reading of this manuscript. The authors also thank the MSU Campus Animal Resources, MSU Mass Spectrometry and Metabolomics Core and the MSU Investigative HistoPathology Laboratory. Funding: This work was supported by the METAvivor Early Career Investigator Grant to SYL.

---

**Supplementary Table 1.**
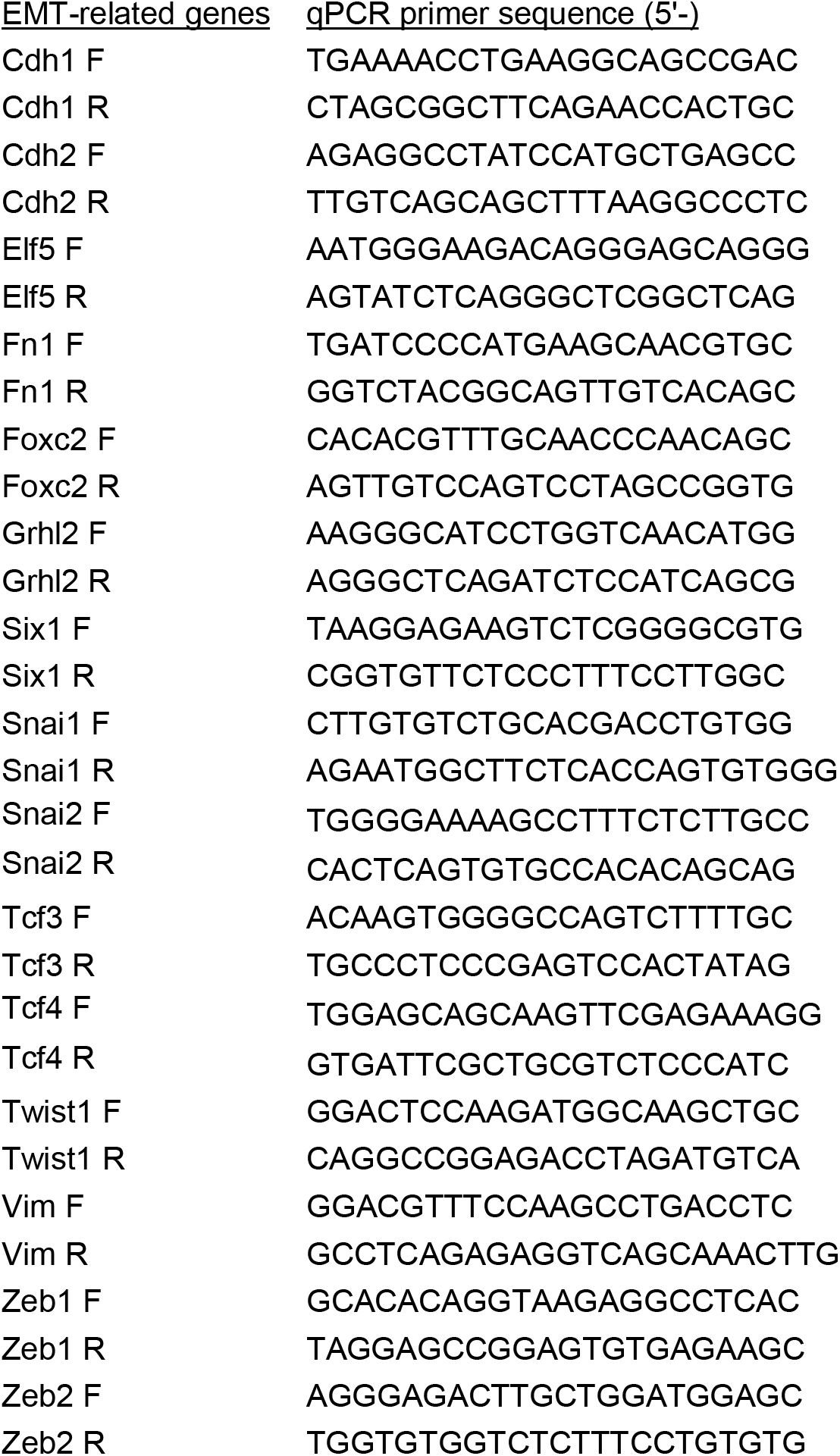
qPCR primers for quantifying expression of EMT-related genes.

**Supplementary Figure 1.**
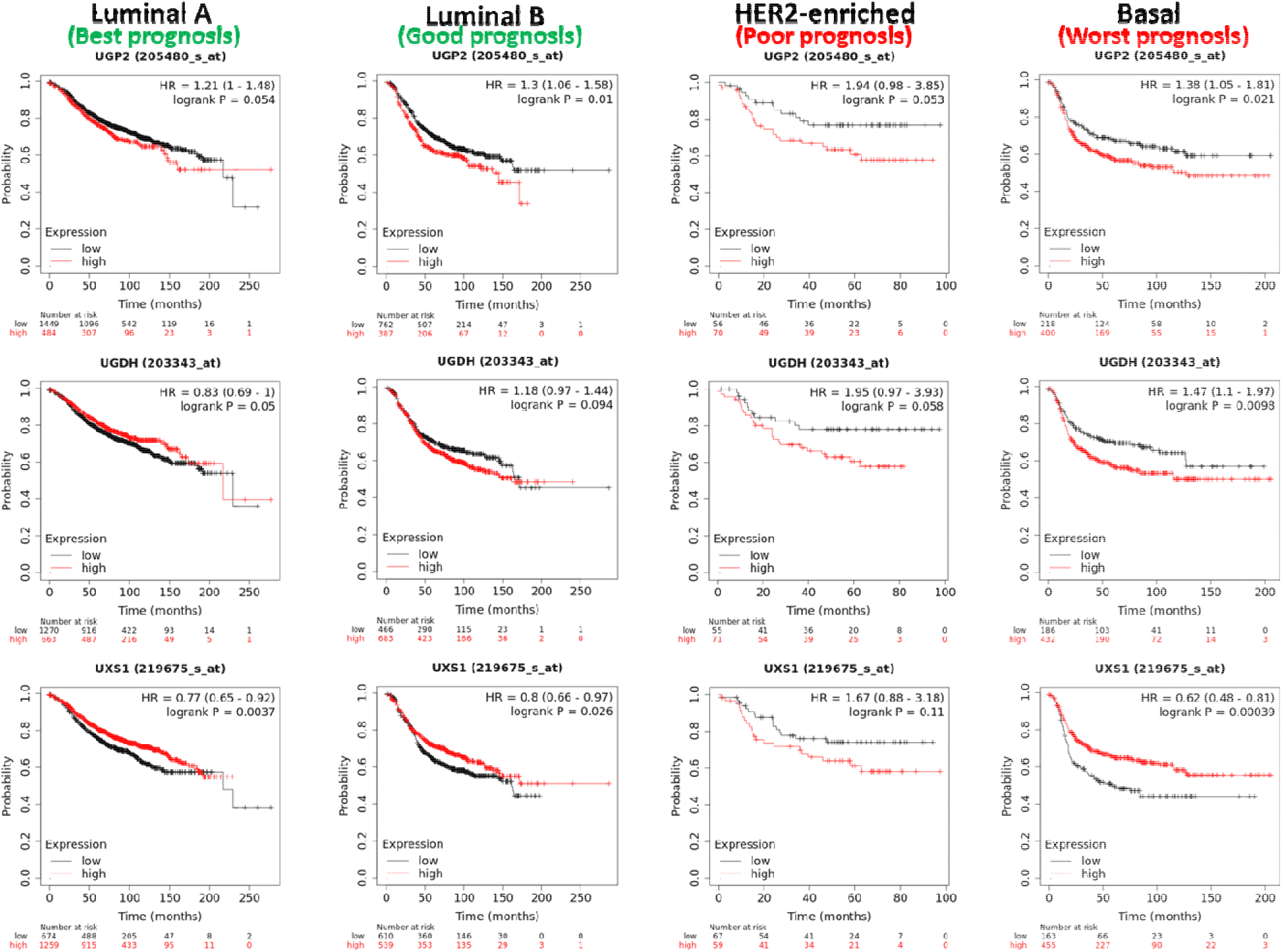
Kaplan-Meier survival curves by breast cancer intrinsic subtype. Kaplan-Meier survival curves comparing relapse-free survival of patients with high versus low expression of UDP-glucose pathway genes, generated separately for patients of each intrinsic subtype. Cut-off values for splitting high-versus low-expression patient groups were automatically determined using the ‘Auto select cutoff’ option in KM Plotter. Hazard ratios and p-values are also listed in **Table 1**.

**Supplementary Figure 2.**
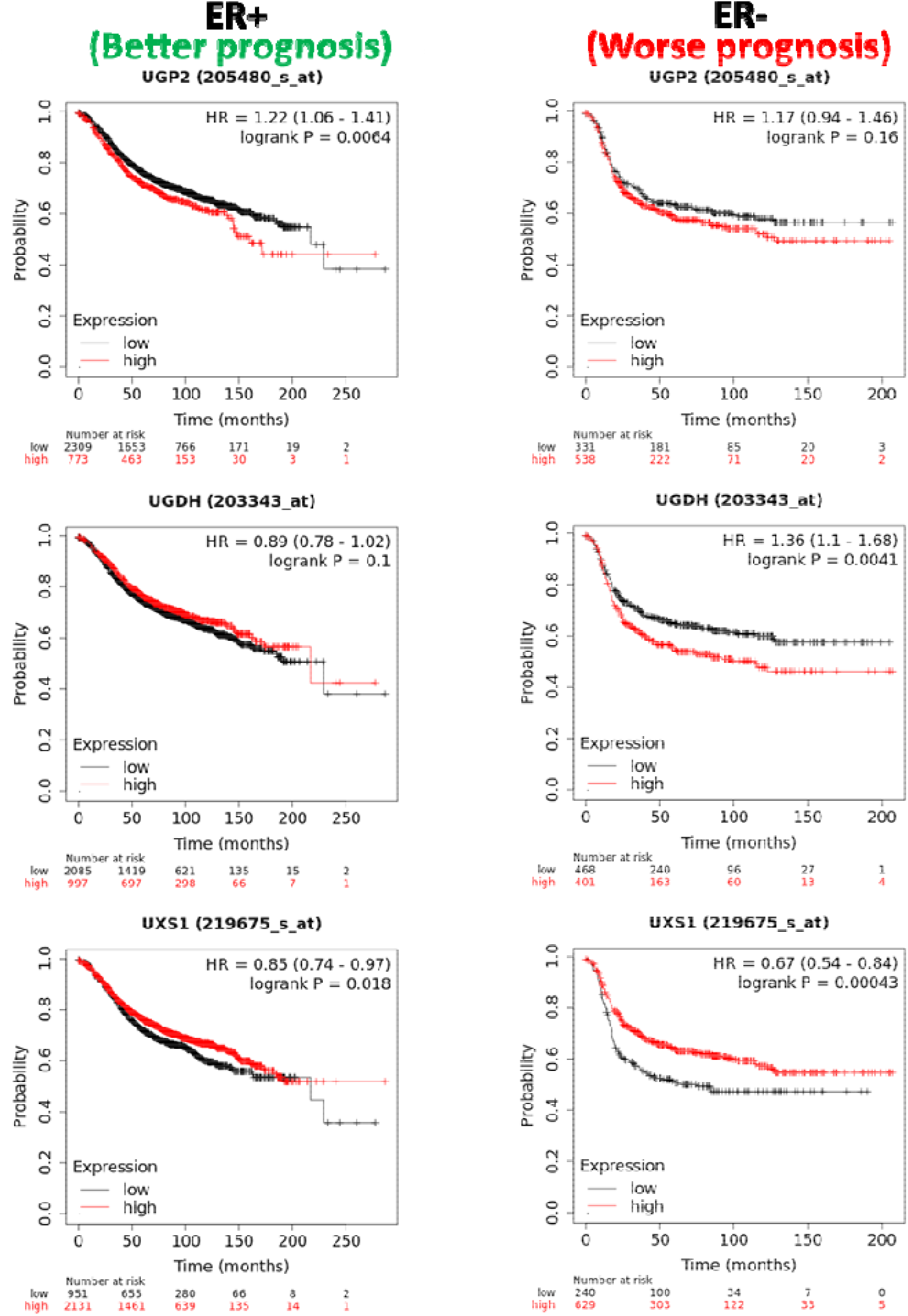
Kaplan-Meier survival curves by estrogen receptor status. Kaplan-Meier survival curves comparing relapse-free survival of patients with high versus low expression of UDP-glucose pathway genes, generated separately for estrogen receptor positive (ER+) and estrogen receptor-negative (ER-) breast cancer patients. Cut-off values for splitting high-versus low-expression patient groups were automatically determined using the ‘Auto select cutoff’ option in KM Plotter. Hazard ratios and p-values are also listed in **Table 1**.

**Supplementary Figure 3:**
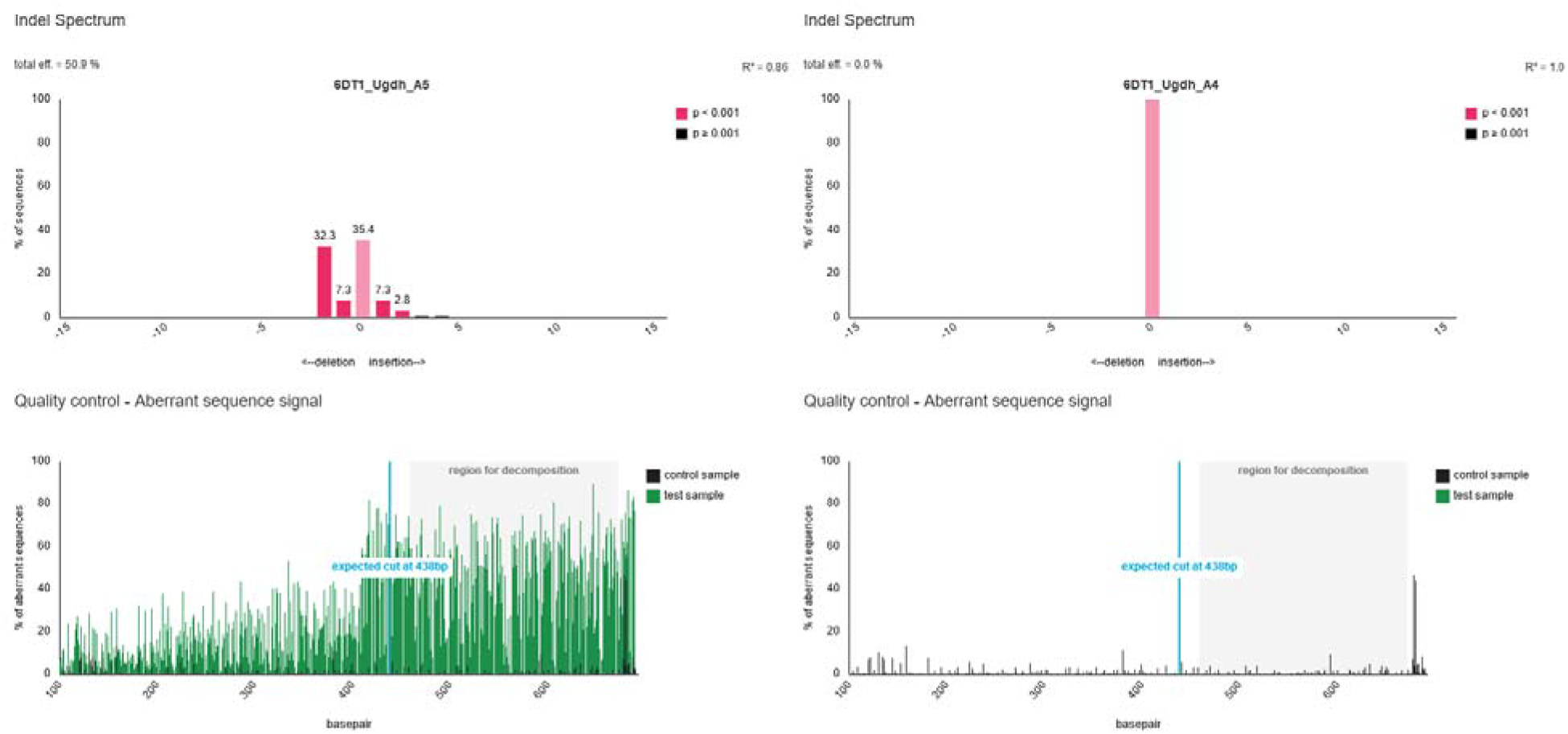
Confirmation of CRISPR/Cas9-mediated Ugdh knockout in 6DT1 cells. Clonal cell populations were examined by TIDE (Tracking of Indels by Decomposition) to verify presence of frame-shift indels in Ugdh DNA sequences. Pink bars denote insertion/deletion events with high confidence (p <0.001). 6DT1_Ugdh_A5 and 6DT1_Ugdh_A4 are used throughout this study as Ugdh-KO and WT control cells, respectively.

**Supplementary Figure 4.**
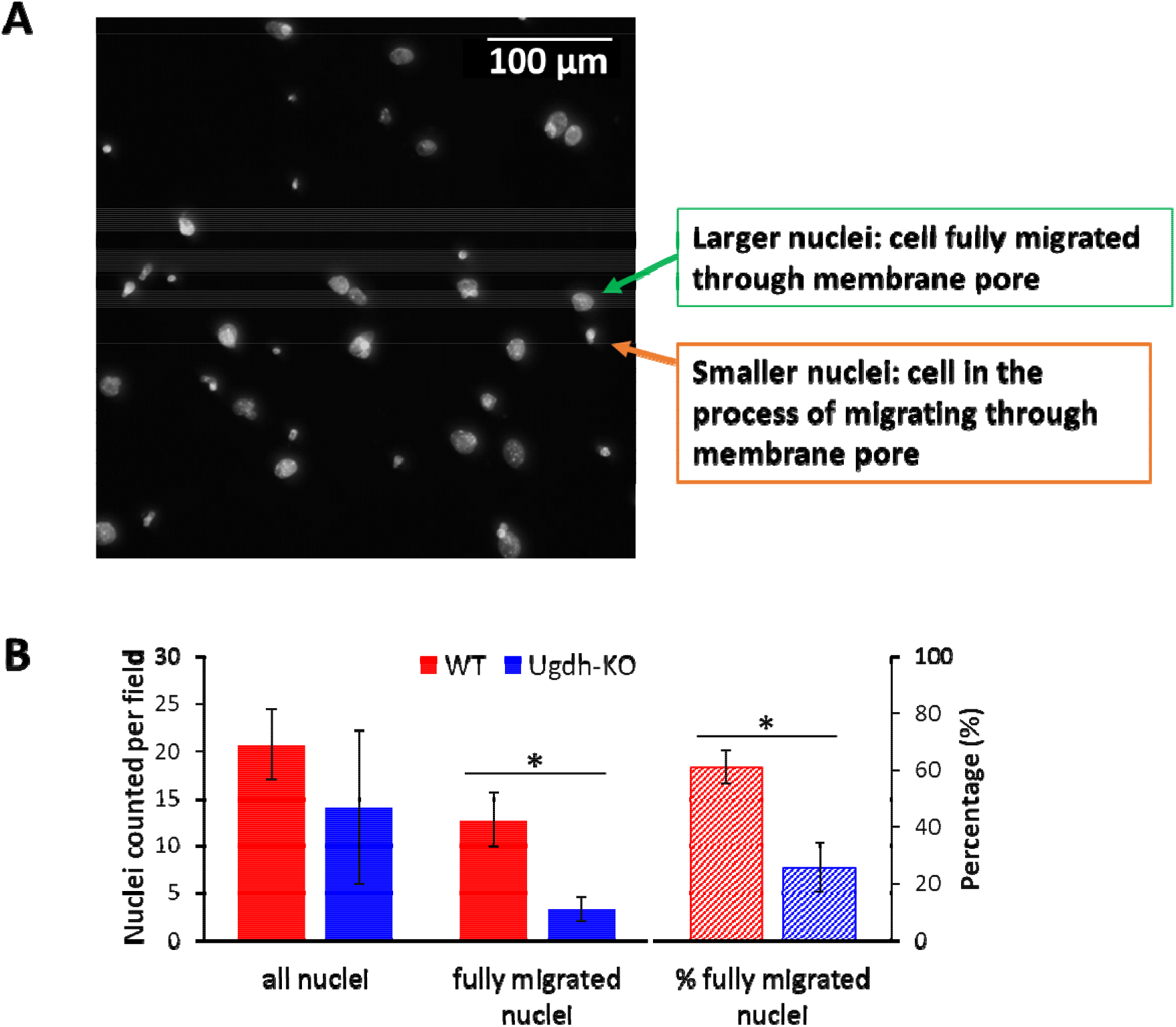
Boyden chamber assay additional details. (A) Representative image of Hoechst 33342 stained nuclei of cells on the bottom surface of Transwell inserts following 16 hours of migration. Fully migrated cells display large, oval-shaped nuclei while partially migrated cells display small, circular nuclei corresponding to the shape and size of the membrane pores. (B) Barplot showing *total* nuclei counted per field, *fully migrated* nuclei counted per field, and percentage of fully migrated nuclei, quantified from images of migrated WT or Ugdh-KO cells (9 images per well, 3 well replicates; total 27 images per cell line). Barplot values and error bars are averages and standard deviations, respectively. Asterisks (*) denote statistically significant difference (p-value < 0.05 by Welch’s t-test).

**Supplementary Figure 5.**
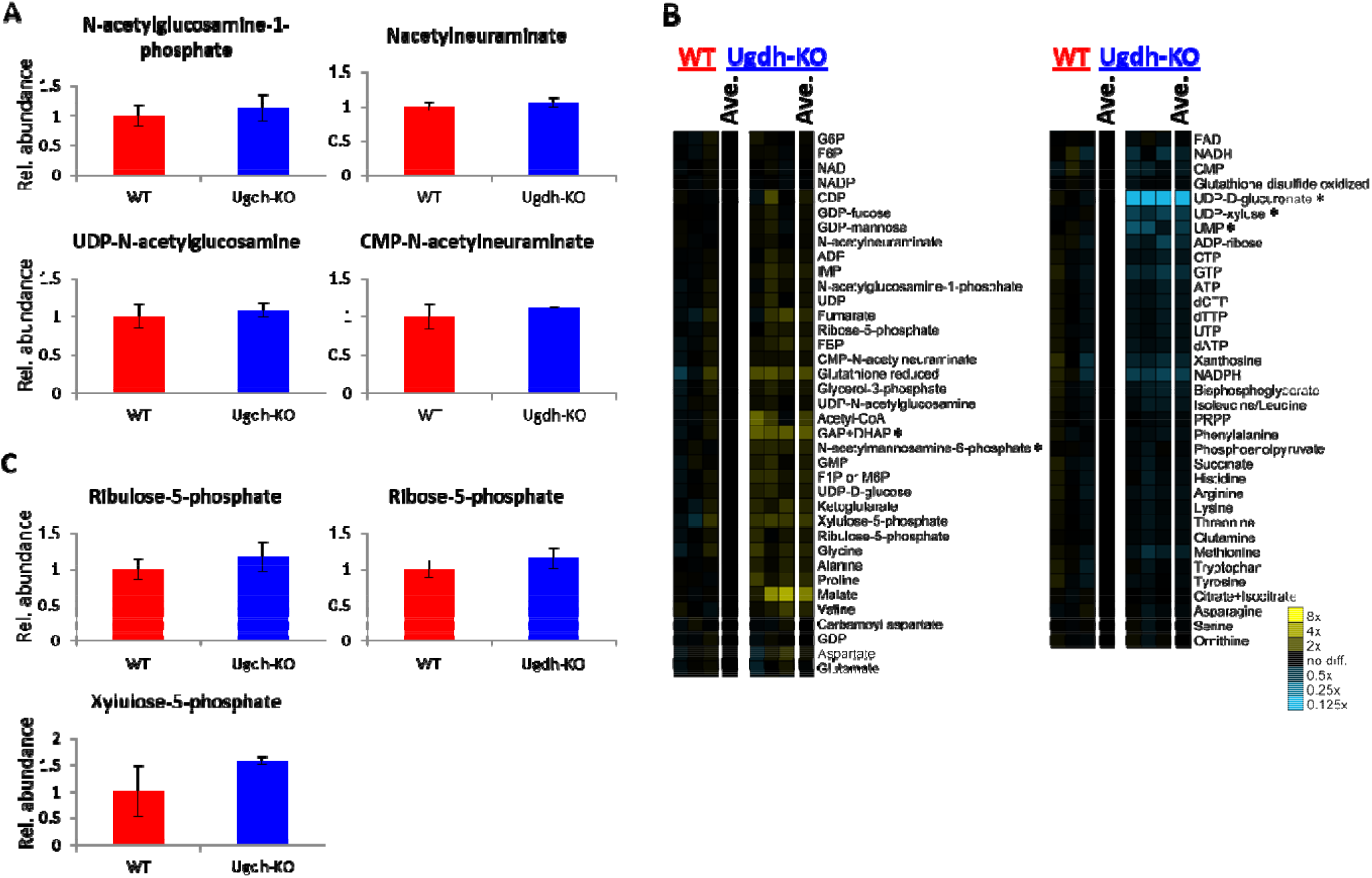
Metabolite profiles of 6DT1 Ugdh-KO vs WT cells. (A) Metabolites in the sialic acid pathway are slightly increased in Ugdh-KO cells. (B) Heatmap showing mostly-similar metabolite profiles in WT and Ugdh-KO cells. Each column represents an individual sample. Yellow and blue cell colors indicate higher and lower metabolite values respectively, relative to average value in WT samples. Metabolite values with statistically significant differences (p-value < 0.05 by Welch’s t test) are marked with asterisks (*). (C) Metabolites in the pentose phosphate pathway are slightly increased in Ugdh-KO cells. Barplot values are the averages of 3 replicates and error bars represent standard deviation.

**Supplementary Figure 6.**
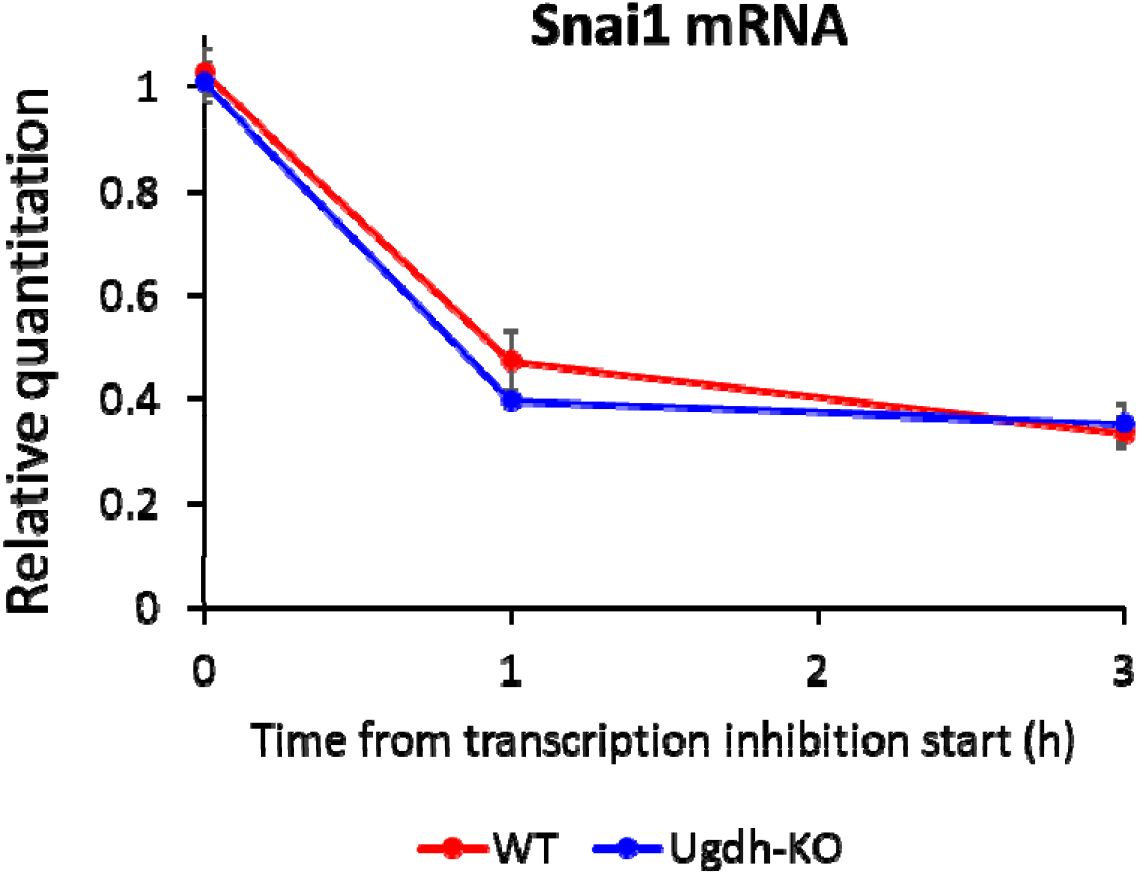
Decreased metastatic ability of Ugdh-KO tumors is not due to differences in *Snai1* expression. Cells were extracted for RNA at indicated time points following transcription inhibition by addition of 1 μg/mL actinomycin D. Snai1 expression levels were measured by qRT-PCR and normalized to control gene Tubb5. Normalized Snai1 expression values are shown relative to t=0 WT samples. Values are averages of three replicates and error bars represent standard deviations. There is no statistically-significant difference between WT and Ugdh-KO at any of the time points.

